# iCOBRA: open, reproducible, standardized and live method benchmarking

**DOI:** 10.1101/033431

**Authors:** Charlotte Soneson, Mark D. Robinson

## Abstract

We present iCOBRA, a flexible general-purpose web-based application and accompanying R package to evaluate, compare and visualize the performance of methods for estimation or classification when ground truth is available. iCOBRA is interactive, can be run locally or remotely and generates customizable, publication-ready graphics. To facilitate open, reproducible and standardized method comparisons, expanding as new innovations are made, we encourage the community to provide benchmark results in a standard format.

---

Many popular tasks undertaken in modern life science research (e.g., in high-throughput studies) can be placed within the broad frameworks of either *ranking* or *classification* of items. For example, differential expression studies can rank genes by the estimated p-value or, using a cutoff on the same, classify genes as either “significantly different” or “not significantly different” between conditions of interest. Other studies combine one or more biomarkers to calculate patient-wise prognostic or predictive risk scores. These can be used to rank the patients in order of, e.g., disease severity, likelihood of treatment response or risk of disease relapse, or to assign samples to different treatment regimes.

Due to the pervasiveness of ranking and classification tasks, a wide range of computational methods have been proposed, often tailored to characteristics of particular data types (e.g., edgeR^1^ or DESeq2^2^ for RNAseq-based differential gene expression, limma^3^ for microarray-based differential gene expression, DEXSeq^4^ for RNAseq-based differential exon usage, metagenomeSeq^5^ for differential abundance analysis of microbiome data and so on). Many of these methods further rely on the accuracy of *estimates* or *quantifications* of underlying entities, such as abundance levels or statistical model parameters. As new methods or variations of existing procedures for these broad classes of computational tasks are developed (classification, estimation or ranking), static comparison or evaluation studies quickly become outdated. Notably, such comparison studies are done as a minimum standard for publishing every new computational method (at least in genomics).

Due to the lack of standard ways to present results from method comparison studies, and the fact that raw results (e.g., simulated data) are not always made available to the readers, it is often difficult for method researchers to reproduce or delve deeper into published evaluations and explore them from different angles, potentially more relevant for specific applications. This stresses the need for open, standardized and reproducible evaluations. Here, we present iCOBRA (interactive Comparative evaluation of Binary classification and RAnking methods), a benchmarking platform for both users and developers of methods. iCOBRA consists of an R package as well as a flexible, interactive web application that can rapidly evaluate and compare methods for binary classification, ranking and continuous target estimation. Methods are evaluated against a ground truth, generally obtained from simulated or independent validation data. In addition, we have collected a set of benchmarking datasets in standard formats, to lower barriers for new method developers but also to facilitate standardized method evaluations in the future.

iCOBRA’s web application is based on the Shiny framework^6^ and implemented in R^7^. It can be run over the web through our public server at http://imlspenticton.uzh.ch:3838/iCOBRA/, which makes it platform-agnostic and eliminates the need for knowledge about installing or running R. Underlying the application is an R package (available through Bioconductor^8^ starting from version 3.3 or from https://github.com/markrobinsonuzh/iCOBRA), which can be used both to run the interactive application locally (e.g., via a web browser) and to generate the result visualizations from the R console directly, facilitating both interactive exploration and integration within programming pipelines. As opposed to other R packages dedicated to evaluating classifiers (e.g., ROCR^9^), which generate static performance plots, the interactivity provided by the Shiny framework lets the user include or exclude methods from a comparison, change the appearance of the plots or stratify the results by a provided annotation with minimal effort. The input format is simple and generic (tab-delimited text with observed results or known truth as columns), leading to an increased ease and range of use compared to other performance evaluators (e.g., compcodeR^10^), whose data representation format and/or choice of evaluation metrics are specifically tailored to certain types of data. The application accepts and harmonizes several types of input values (nominal p-values, adjusted p-values and a general “score”), allowing for flexibility beyond existing applications like BDTcomparator^11^, which compares two categorizations and is thus strictly limited to classification evaluation. Depending on the provided inputs, iCOBRA will select the optimal one to use for calculation of each evaluation metric. Finally, the flexibility of iCOBRA allows the results for different methods to be uploaded either together or separately, and missing predictions can be handled according to the user’s wish.

Figure 1 shows a screenshot of the iCOBRA web interface. The left panel contains the input controls, while the tabs along the top show different evaluation panels. The inputs provided by the user are a “truth” file, containing the ground truth for each considered item as a binary or continuous variable, and one or more “result” files, providing one or more of the three types of inputs (p-value, adjusted p-value or score) for each item and each evaluated method. The truth file can also contain other annotations that can be used to stratify the evaluation. In Figure 1, the visualization of the results has been stratified into three groups by the annotation called “diff_rel_abundance”, which in this case represents the true relative abundance difference between differentially used isoforms. The method performances for each of the three groups of genes are shown in separate panels alongside the overall performance.

**Figure 1.**
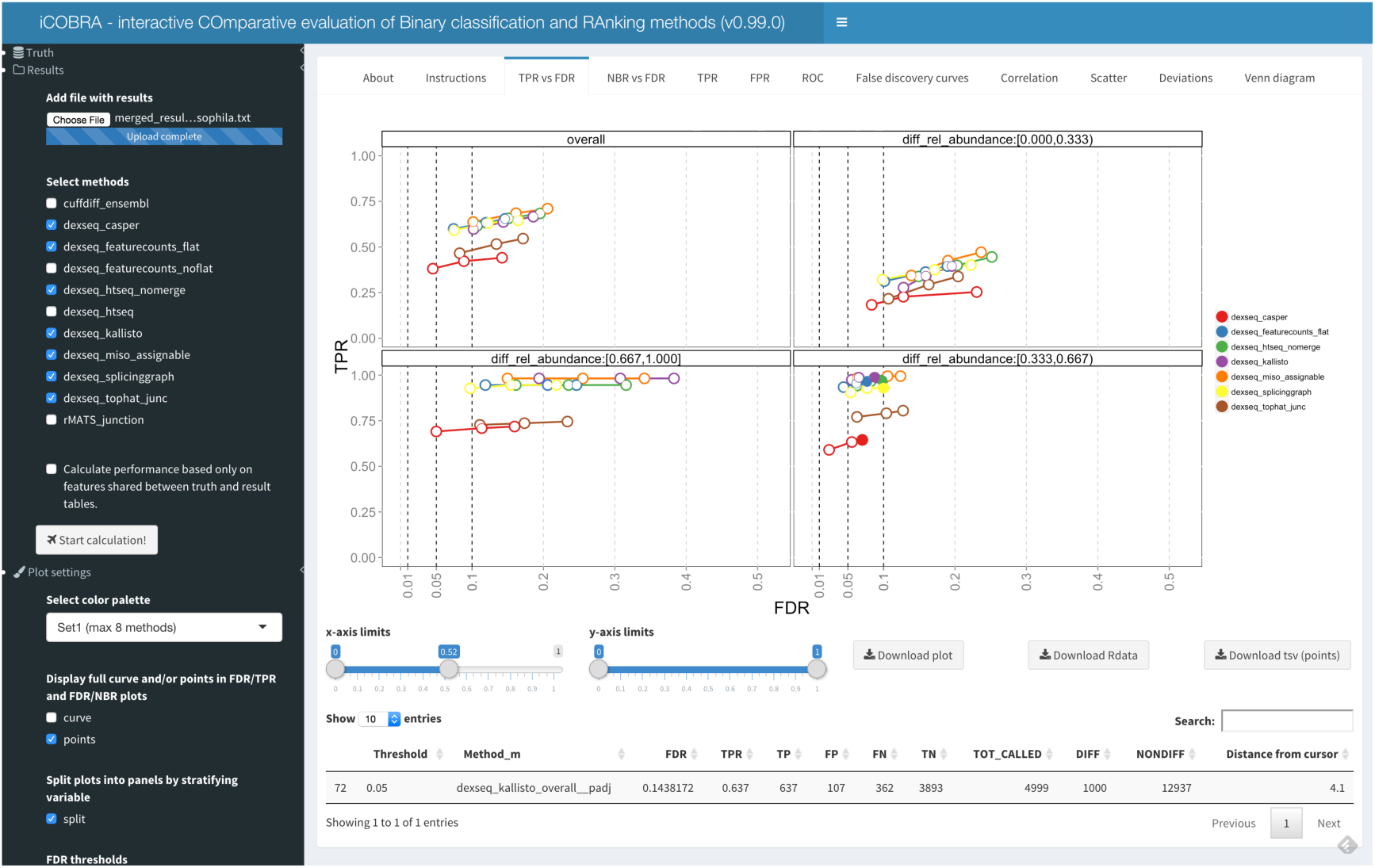
Screenshot of the iCOBRA interactive application interface. The left sidebar contains the input controls, while the tabs in the main panel display different aspects of the performance evaluation.

The displayed figure shows the observed true positive rate (TPR, power, sensitivity, recall) versus the observed false discovery rate (FDR) at three (customizable) adjusted p-value cutoffs, with different methods represented by different colors. By checking the box denoted “curve” in the input panel to the left, the entire TPR-vs-FDR curve (traced out by going through all possible cutoff values) can also be displayed. This is especially useful in situations where only a general score is available for one or several methods. If the cursor is hovered over the figure, the table beneath the plot displays the coordinates and other information for the points in the vicinity of the cursor. These graphics can also be created by loading the truth and result files into an R session and using the object classes and functions provided within the iCOBRA package.

Other commonly-used evaluation metrics available in iCOBRA include “NBR vs FDR” (total number of items classified as positive against the achieved FDR), “TPR” (achieved true positive rate at selected adjusted p-value cutoffs), “FPR” (achieved false positive rate at selected adjusted p-value cutoffs), “ROC” (receiver operating characteristic curves), “false discovery curves” (number of items falsely classified as positive among the top-ranked items), “correlation” (correlation between provided scores and a continuous target), “scatter” (display of observed and true scores), “deviations” (distribution of deviations between observed and true scores) and “Venn diagram” (overlap among the sets of items called positive by the different methods). All plots can be exported in pdf format, and the underlying data can be exported in RData format or as tab-separated values for further exploration, using the iCOBRA R package or other tools.

We believe that the interactivity and ease of use of iCOBRA makes it a valuable tool for general comparison and evaluation of binary classification, ranking and estimation methods, and that it can help to drive benchmarking efforts forward as the method toolbox is extended. Authors of new methods and comparison studies can easily publish the two text files that make it possible for any reader to, within minutes, reproduce and further explore and extend the published evaluations. Of course, researchers are also encouraged to make full details of the primary data going into the comparison available in order to facilitate full reproducibility. In addition, the iCOBRA package and web application are extensible, if additional performance metrics and visualizations are preferred.

We have collected several benchmarks containing recent example datasets suitable for evaluation of methods for detection of differential gene or transcript expression^13^, differential transcript usage^13^, transcript abundance estimation^14^, relative inclusion of alternative splicing events^15^ and metagenome abundance estimation (http://imlspenticton.uzh.ch/robinsonlab/benchmarkcollection/). These datasets can be used as a basis for standardized method evaluation in these application areas and include direct links to the original data as well as typical intermediate file formats along standard pipelines. We envision this database to be extended over time and encourage the community to contribute their method assessments to the public domain.

